# Deeply conserved super-enhancers maintain stem cell pluripotency in placental mammals

**DOI:** 10.1101/2022.05.03.490430

**Authors:** Juqing Zhang, Yaqi Zhou, Wei Yue, Zhenshuo Zhu, Xiaolong Wu, Shuai Yu, Qiaoyan Shen, Qin Pan, Wenjing Xu, Rui Zhang, Xiaojie Wu, Xinmei Li, Yayu Li, Yunxiang Li, Yu Wang, Sha Peng, Shiqiang Zhang, Anmin Lei, Xinbao Ding, Fan Yang, Xingqi Chen, Na Li, Mingzhi Liao, Wei Wang, Jinlian Hua

**Affiliations:** College of Veterinary Medicine, Shaanxi Centre of Stem Cells Engineering & Technology, Northwest A&F University, Yangling, Shaanxi 712100, China; National Institute of Biological Sciences, Beijing, 102206, China; College of Life Sciences, Northwest A&F University, Yangling, Shaanxi, 712100, China; College of Animal Sciences& Technology, Northwest A&F University, Yangling, Shaanxi 712100 China; Cornell University, College of Veterinary Medicine, Department of Biomedical Sciences, Ithaca, NY 14853, USA; Department of Cell and Molecular Biology, Karolinska Institutet, Solna 17177, Sweden; Tsinghua Institute of Multidisciplinary Biomedical Research, Tsinghua University, Beijing, China

**Keywords:** Pluripotency, Super-enhancer, Evolution, Pigs, BRD4

## Abstract

Despite pluripotent stem cells sharing key transcription factors, their maintenance involves distinct genetic inputs. Emerging evidence suggests that super-enhancers (SEs) can function as master regulatory hubs to control cell identity and pluripotency in humans and mice. However, whether pluripotency-associated SEs share a deep evolutionary origin in mammals remains elusive. Here, we performed comprehensive comparative epigenomic and transcription factor binding analyses among pigs, humans, and mice to identify pluripotency-associated SEs. Like typical enhancers, SEs displayed rapid evolution in mammals. We showed that BRD4 is an essential and conserved activator for mammalian pluripotency-associated SEs. Comparative motif enrichment analysis revealed 30 shared transcription factor binding motifs among the three species. The majority of the transcriptional factors that bind to identified motifs are known regulators associated with pluripotency. Further, we discovered three pluripotency-associated SEs (SE-SOX2, SE-PIM1, and SE-FGFR1) which displayed deep conservation in placental mammals and are sufficient to drive reporter gene expression in a pluripotency-dependent manner. Disruption of these conserved SEs through the CRISPR/Cas9 approach severely impaired the proliferative potential and the ability to form undifferentiated colonies. Our study provides insights into the understanding of conserved regulatory mechanisms underlying the maintenance of pluripotency as well as species-specific modulation of the pluripotency-associated regulatory networks in mammals.

**Significance statement:** Super-enhancers (SEs) hold stronger power than regular enhancers to direct gene expression in the regulation of stem cell pluripotency. To dissect how pluripotency-associated SEs have evolved in mammals, we performed a systematic comparison of SEs among pigs, humans, and mice. Our analysis allowed the identification of three pluripotency-associated SEs (SE-SOX2, SE-PIM1, and SE-FGFR1) that are highly conserved in *Placentalia* (accounting for 94% of mammals) as well as many species-specific SEs. All three SEs were sufficient to direct pluripotency-dependent gene expression and disruption of each conserved SE caused the loss of stem cell pluripotency. Our work highlights a small number of highly conserved SEs essential for the maintenance of pluripotency.

## Introduction

The developmental potential of interspecific chimeras established from pluripotent stem cells (PSCs) indicates the presence of conserved regulatory mechanisms controlling the proliferation and differentiation during development (1, 2). Mammalian PSCs share key transcription factors and can be induced from adult somatic cells through the forced expression of OCT4, SOX2, KLF4, and c-Myc to activate pluripotency-associated genes (3-5). However, PSCs derived from different species require distinct external signals, suggesting a regulatory discrepancy in maintaining the pluripotency among different species (6, 7). For instance, MEK-ERK inhibitors (such as PD-0325901) have an opposing effect on boosting pig (*Sus scrofa*) and mouse pluripotency (8). How the regulatory landscape responsible for the maintenance of stem cell pluripotency has evolved in mammals is still unclear.

Enhancers or *cis*-regulatory elements play key roles in controlling spatial and temporal expression of genes essential for many biological processes (9). Emerging evidence suggests that groups of enhancers with unusually strong enrichment for the binding of transcriptional coactivators may accumulate at certain regions of animal genomes to form super-enhancers (SEs) (10). Compared with typical enhancers (TEs), SEs are characterized by high levels of histone H3 lysine 27 acetylation (H3K27ac) density with an enormous size (11, 12). Such SEs can function as master regulatory hubs to determine cell identity, developmental and disease states in a broad range of human and mouse cell types (13-15). Previous studies have shown that SEs in mouse PSCs are densely occupied by master pluripotency-related transcription factors and reduced expression in OCT4 or mediators led to preferential loss of expression of SE-associated genes relative to other genes (16). Deletion of the SEs associated with NANOG or SOX2 in mice blocked its corresponding gene expression and caused a failure in the maintenance of pluripotency (17-20).

Since the majority of studies on mammalian pluripotency were conducted in humans and mice (21-23), whether pluripotency-associated SEs share a deep evolutionary origin in mammals is still an open question in the field (24, 25). To address this question, we extended our research to pigs that are distantly related to humans and mice. Pigs have been used as a critical model for numerous human conditions and diseases including xenotransplantation due to the similarity in organ anatomy, physiology, and metabolism (26, 27). For instance, genome-edited pigs were reported to better recapitulate the features of neurodegenerative diseases compared to similarly modified mice models (28). Recently, several porcine pluripotent stem cell lines derived from preimplantation embryos or established through induced pluripotent stem cell technology showed the ability to differentiate into the three germ layers (29, 30). These PSC lines provide a unique opportunity for investigating the evolution of SEs associated with pluripotency in mammals. In this study, we performed systematic identification of SEs in porcine PSCs using multi-omics. Our cross-species comparative epigenomic analysis revealed three highly conserved SEs essential for stem cell pluripotency in mammals.

## Results

### Systematic identification of super-enhancers associated with porcine pluripotent stem cells

Reprogramming porcine somatic cells by doxycycline (Dox) induced expression of four transcriptional factors (OCT4, SOX2, KLF4, and c-Myc) is an efficient approach to produce porcine pluripotent stem cells (PSCs) (8, 31, 32). Using this system, we have established a porcine pluripotent cell line possessing the ability to produce xenochimerism with mouse early embryos (33) (Fig. 1A and B). Changes in culture condition are sufficient to drive these porcine PSCs to switch rapidly between pluripotent state and differentiation state, as indicated by alkaline phosphates (AP) staining and the expression levels of pluripotent genes (Fig. 1C and D). Therefore, we chose this cell line for the identification and characterization of SEs in this study.

**Figure 1.**
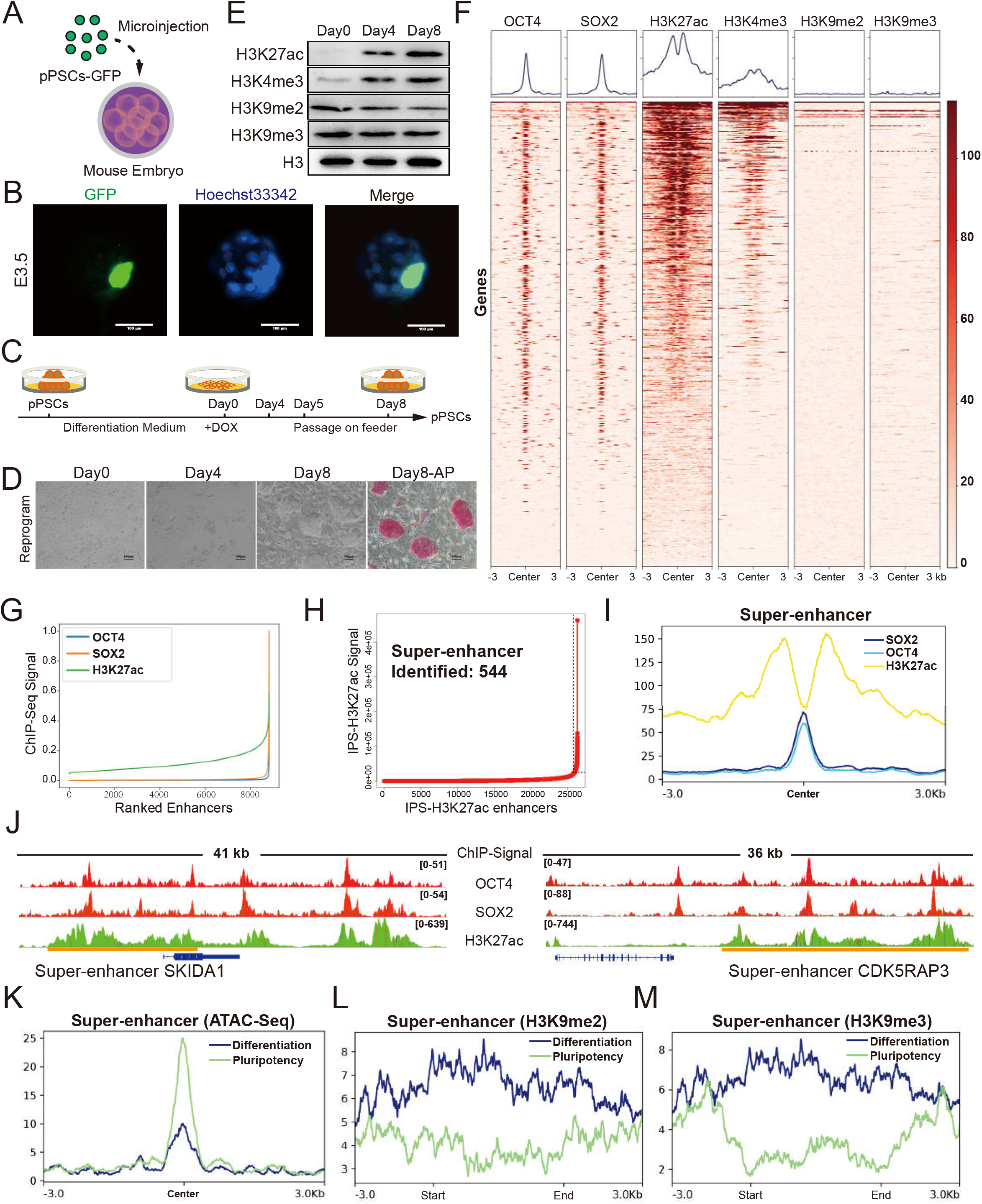
Systematic identification of super-enhancers in porcine pluripotent stem cells. *(A)* Schematic of the generation of pig-mouse *ex-vivo* chimeric embryos. *(B)* Representative images of porcine PSC-injected mouse embryos cultured *in vitro* at E3.5. The nuclei were marked by hoechst33342 and the location of the injected porcine PSCs was marked with GFP. Scale bar, 100 µm. *(C, D)* Strategies for switching porcine PSCs between pluripotent state and differentiation state by changing the culture condition. Under the reprogramming culture condition, differentiated porcine PSCs transformed to the pluripotent state (as referred to secondary reprogramming). Scale bar, 100 µm. *(E)* Western blot analysis of representative histone modifications during secondary reprogramming. *(F)* Heatmaps of OCT4, SOX2, H3K27ac, H3K4me3, H3K9me2, and H3K9me3, surrounding the OCT4 binding loci. *(G)* Distribution of H3K27ac, OCT4, and SOX2 normalized ChIP-Seq signals. Each ChIP-seq data point was normalized by dividing the ChIP-seq signals by the maximum signals individually and sorted in ascending order. *(H)* Identification of SEs in porcine PSCs. Within the 12.5-kb window, H3K27ac signals were ranked for porcine PSCs. The enhancers above the inflection point were defined as original super-enhancers. *(I)* Average ChIP-seq signal density of OCT4, SOX2, and H3K27ac surrounding the center of the SEs regions. *(J)* ChIP-seq signal profiles of OCT4, SOX2, and H3K27ac in porcine PSCs at two representatives top-ranked SEs locus. Gene models were shown below the binding profiles. The positions of SEs were marked by orange lines. *(K)* Average ATAC-seq signal density in pluripotent state and differentiation state of porcine PSCs surrounding the SEs regions. *(L, M)* Average ChIP-seq signal density of H3K9me2 and H3K9me3 in pluripotent and differentiation state of porcine PSCs surrounding the SEs regions.

H3K27ac histone modification, combined with others such as H3K4me3 (marks active promoters), has been frequently used as a marker for the identification of active TEs and SEs (34-36). Previously, SEs associated with stem cells were defined by the presence of sites bound by master regulators (such as OCT4, SOX2, and NANOG), and by the potential to span large genomic regions, with an average size generally an order of magnitude larger than that of TEs (14, 16, 37). Upon the establishment of pluripotency in PSCs, the levels of total H3K27ac and H3K4me3 histone modification increased dramatically (Fig. 1E). In contrast, the repressive histone marks H3K9me2 and H3K9me3 displayed a slight but statistically significant reduction (Fig. 1E). To identify SEs associated with porcine PSCs, we performed chromatin immunoprecipitation sequencing (ChIP-seq) including histone ChIP-seq (H3K27ac, H3K4me3, H3K9me2, and H3K9me3) and transcription factor ChIP-seq (OCT4 and SOX2). Our results showed that 2159 out of 3337 OCT4- and SOX2-regulated genes displayed H3K27ac enrichment (Fig. 1F). Consistent with previously reported observations in humans and mice, OCT4- and SOX2-enriched peaks overlapped well with H3K27ac-defined enhancers in pigs (Fig. 1G). Next, SEs associated with porcine PSCs were identified by concatenated H3K27ac-marked regions (absent for H3K4me3 signals), the presence of OCT4 and SOX2 ChIP signals, and a minimum size of 5.8 kb (38) (Fig. 1H and I). For instance, we found there were two top-ranked SEs associated with SKI/DACH Domain Containing 1 (SKIDA1), and CDK5 Regulatory Subunit Associated Protein 3 (CDK5RAP3), respectively (Fig. 1J). The latter has been reported to be essential for the survival of human PSCs (39). GO term analysis suggested that the identified SE-associated genes are associated with embryo development and signaling pathways regulating the pluripotency of stem cells. To further confirm the identified SEs are indeed associated with pluripotency, we conducted ATAC-seq to monitor the chromatin accessibility in SE regions in both pluripotent state and differentiation state. We found that SE regions were open in the pluripotent state, but exhibited a dramatic reduction in chromatin accessibility during the differentiation of porcine PSCs (Fig. 1K). Consistently, SE regions were strongly marked by repressive histone marks H3K9me2 and H3K9me3 in the differentiation state but displayed relatively weak signals in the pluripotent state (Fig. 1L and M). Taken together, we concluded that the SEs we have identified in porcine PSCs are closely associated with pluripotency.

### BRD4 is an essential activator for mammalian pluripotency-associated SEs

BRD4, a highly conserved (> 84% sequence identity between humans and basal mammals) chromatin reader protein that recognizes and binds acetylated histones, directly interacts with OCT4 and SOX2 to activate pluripotency-related genes in mice (40-42) (Fig. 2A). To investigate whether this regulatory mechanism is common in mammalian PSCs, we performed BRD4 ChIP-seq and compared the BRD4 enrichment in SEs identified from pigs, humans, and mice. We observed a consistent enrichment of BRD4 binding in SEs identified from all three species (Fig. 2B). Co-immunoprecipitation experiments supported that the direct interaction between BRD4 and OCT4 was also present in porcine PSCs (Fig. 2C). To further explore the function of BRD4 in porcine PSCs, we attempted to establish a BRD4-KO cell line using the CRISPR/Cas9 approach. A GFP-STOP cassette was introduced to produce a BRD4 mutant allele, while another RFP-STOP cassette was used to destroy the second allele to produce BRD4 homozygous knockout cells. However, no GFP/RFP double-positive cell was obtained, providing direct evidence for the necessity of BRD4. Therefore, we switched to BRD4 inhibitors and RNAi. We found BRD4 inhibition through drug treatment (JQ1 and iBET762) led to rapid differentiation of porcine PSCs (Fig. 2D), as revealed by EdU assays and cell growth curves. Similarly, BRD4-knockdown blocked the establishment of porcine PSCs, resulting in a failure in the formation of typical porcine PSC colonies (Fig. 2E). Additionally, the phenotype of BRD4 knockdown seems to be more robust compared with the knockdown of E1A binding protein P300 (an important histone acetylase that is extensively involved in SEs-associated transcriptional regulation) (Fig. 2E).

**Figure 2.**
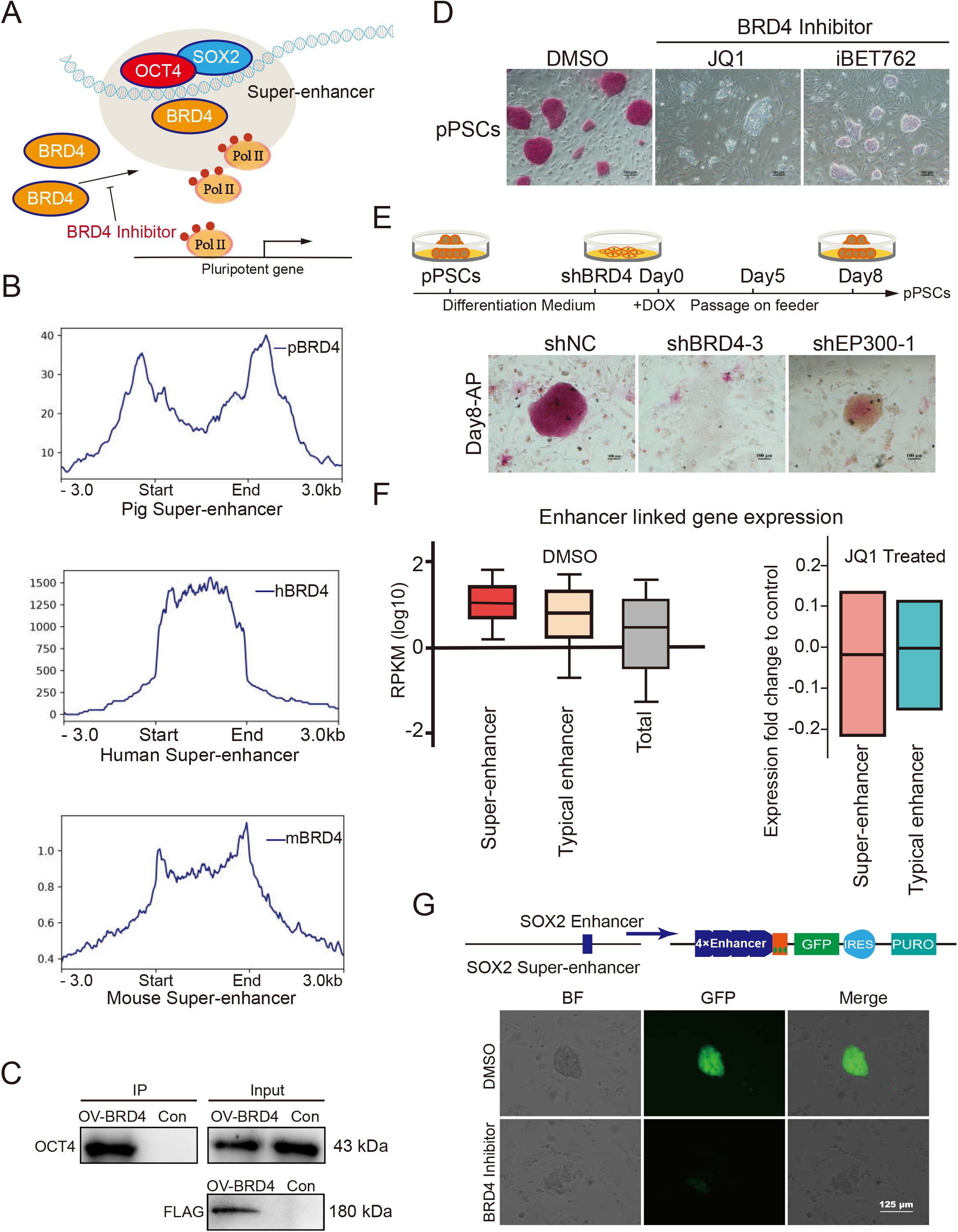
BRD4 is an essential activator for mammalian pluripotency-associated SEs. *(A)* Model showing pluripotency-associated SEs working relying on the enrichment of BRD4 in human and mouse PSCs. *(B)* Average ChIP-seq signal density of BRD4 in pigs, humans, and mice surrounding the pluripotency-associated SEs regions. *(C)* The interaction of BRD4 and OCT4 in porcine PSCs. FLAG-tagged BRD4 was revealed by Western blot using the anti-Flag antibody. We used the anti-OCT4 antibody to test the protein interaction in porcine PSCs. *(D)* AP staining assay of porcine PSCs treated with iBET-762, JQ1, and DMSO. Porcine PSCs lost their pluripotency after BRD4 inhibitor (iBET-762 and JQ1) treatment. Scale bar, 100 µm. *(E)* The strategy for reprogramming BRD4-inhibited porcine cells. The top part is the schematic to reprogram the differentiated PSCs with BRD4 interference. The AP staining assay for cells carrying different shRNAs after 8 days of reprogramming is shown at the bottom. *(F)* Box plots of expression (reads per kilobase of exon per million mapped reads (RPKM)) from enhancer-linked genes and all expressed genes. Genes linked to SEs have relatively high expression levels and inhibition of BRD4 leads to a slightly greater reduction in the expression of SE-linked genes compared with that of TE-linked genes. *(G)* GFP expression driven by the SOX2 enhancer in mouse PSCs. All images were based on the same exposure conditions. The BRD4 inhibitor reduced the activity of super-enhancers. Scale bar, 125 µm.

Then we wanted to ask if the phenotype we observed for BRD4 inhibition is correlated with the reduction in enhancer activities. Therefore, we performed a unique identifier transcriptome profiling of JQ1-treated porcine PSCs. We found BRD4 inhibition led to the suppression of pluripotency-related signaling pathways. Notably, we observed that the expression of genes linked to SEs is more robust than that of TEs in porcine PSCs (Fig. 2F). Blocking BRD4 led to a slightly greater reduction in the expression of SE-linked genes compared with TE-linked genes (Fig. 2F). Next, we tested if BRD4 is essential for the activation of SEs in pigs. Using the SE associated with SOX2 (a known regulator of pluripotency) as an example, we found that inhibition of BRD4 blocked the cell pluripotency and the activation of SOX2 SE in a reporter assay (43) (Fig. 2G). In summary, our data suggest BRD4 is a highly conserved regulator essential for the activation of SEs in mammals.

### Pluripotency-associated transcription factor binding motifs are enriched in SEs

We next explored the motif enrichment of pluripotency-associated SEs in pigs, mice, and humans. Our analysis revealed 278 significantly enriched transcription factor binding motifs for humans SEs, 345 for pigs, and 302 for mice (Fig. 3A). Among these predicted transcription factors, 191 of them were shared in all three species (Fig. 3B). Combined with the transcriptome data of humans, pigs, and mics PSCs, we identified 30 shared pluripotency-related transcription factors with relatively high abundance. To test whether these predicated transcription factors indeed bind to SEs, we took GABPA as an example and examined its binding regions in genomes of human and mouse PSCs using previously published ChIP-seq datasets (44, 45). As expected, we observed significant enrichment of GABPA occupation in SEs regions in both humans and mice (Fig. 3C). Protein-protein interaction analysis using Cytoscape revealed that complex interactions are likely to present among these 30 transcription factors (Fig. 3D).

**Figure 3.**
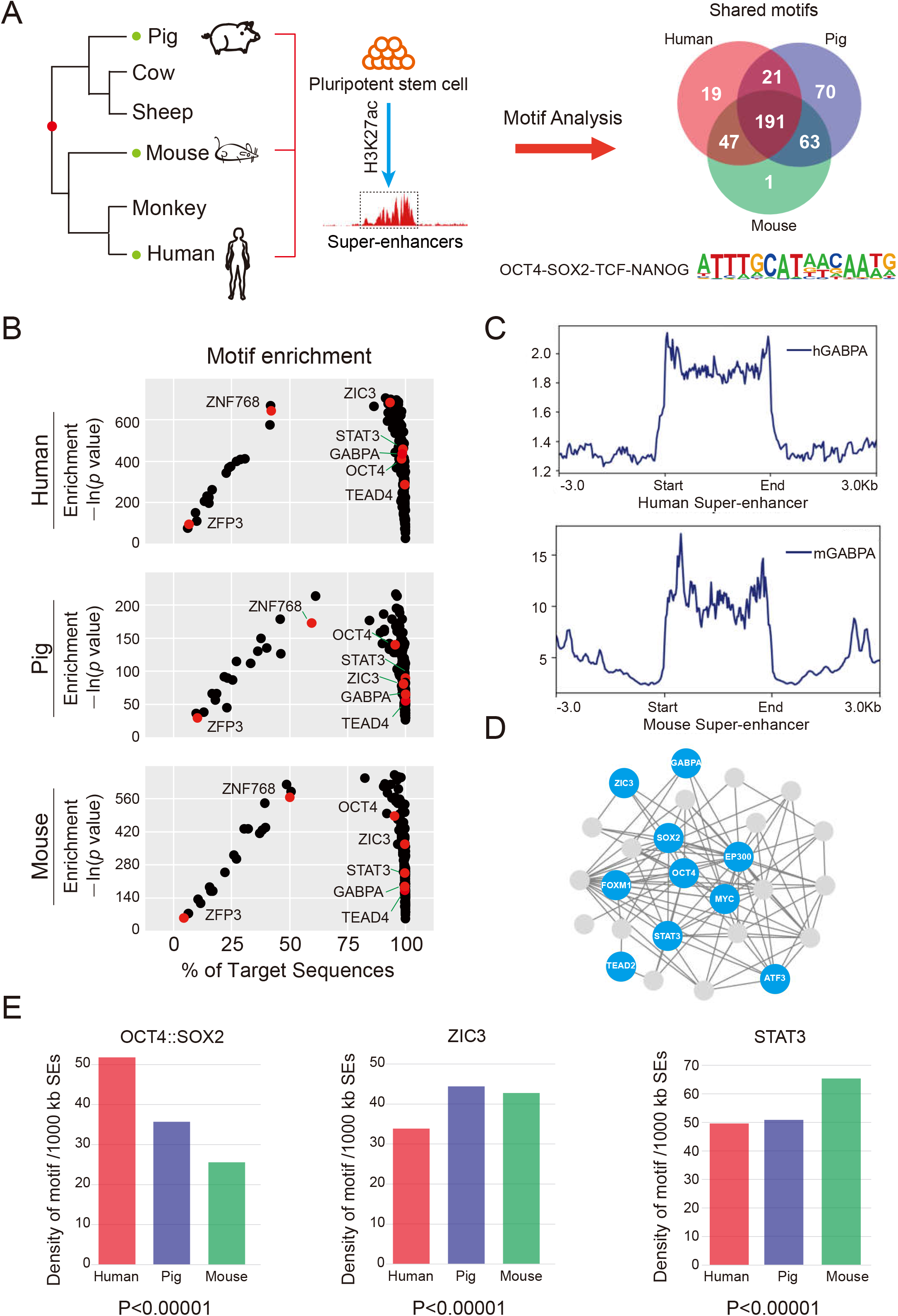
Pluripotency-associated transcription factor binding motifs are enriched in SEs. *(A)* The evolution relationship between humans, pigs, and mice. SEs sequences from the three species were extracted for motif analysis. *(B)* The distribution of the 191 overlapped motifs identified in human (top), pig (middle), and mouse (bottom) pluripotency-associated SEs. Representative pluripotency-associated factors were marked in red. *(C)* Average ChIP-seq signal density of GABPA in humans and mice pluripotency-associated SEs. GABPA is enriched in SE regions. *(D)* Protein interaction networks of representative transcription factors in pigs. Factors with relatively high expression levels were marked in blue. *(E)* The average numbers of the OCT4::SOX2 motif, the ZIC3 motif, and the STAT3 motif per 1000 kb SEs.

Further, the genome-wide distribution of the representative motifs (*e*.*g*., OCT4::SOX2, ZIC3, and STAT3) was compared among pigs, mice, and humans. We observed a large variation of the relative density of each binding motif in different species (Fig. 3E). For example, the density of the OCT4::SOX2 motif in human SEs is about twice higher than that of mice. To overcome the bias caused by species-specific modulation of motifs, we also calculated the motif distribution with human- and mouse-specific motif sequences and observed consistent trends in the tested species. Taken together, our data indicate conserved transcription factor binding motifs are present but not evenly distributed in mammalian SEs associated with stem cell pluripotency, which may explain the diversity in the regulatory landscape of pluripotency.

### Highly conserved pluripotency-associated SEs are present in mammals

Unlike protein-coding genes, enhancers or *cis*-regulatory elements usually exhibit weak DNA sequence conservation and undergo rapid evolution in mammals (46, 47). Nevertheless, certain ancient groups of highly conserved enhancers have been identified in vertebrate development (48, 49). For instance, a limb-specific enhancer of the Sonic hedgehog, the zone of polarizing activity regulatory sequence (ZRS), is highly conserved in many vertebrate species (50). Genomic substitution of the mouse enhancer with its snake orthologs caused severe limb reduction, suggesting that highly conserved enhancers are functionally significant. To determine whether highly conserved pluripotency-associated SEs are present in mammals, we compared the distribution and conservation of pluripotency-associated SEs and TEs in the three distantly related mammals. Our results unveiled 1409 SEs and 23797 TEs in humans, 1108 SEs and 17277 TEs in mice, and 544 SEs and 25786 TEs in pigs (Fig. 4A). To understand the functions of identified enhancers, we combined transcriptome data, H3K4me3 (marks active promoters) ChIP-seq data, and Hi-C data (51, 52), to link enhancers to their potential target genes. We identified 3067 shared TE-linked genes (about 27.7 % of porcine TE linked genes) and 58 shared (about 11.2 % of total porcine SE linked genes) SE-linked genes in all three species (Fig. 4B). GO term analysis indicated that the top GO term associated with shared SE-linked genes and TEs-linked genes was regulation of cell proliferation and establishment of cell polarity, respectively (Fig. 4C).

**Figure 4.**
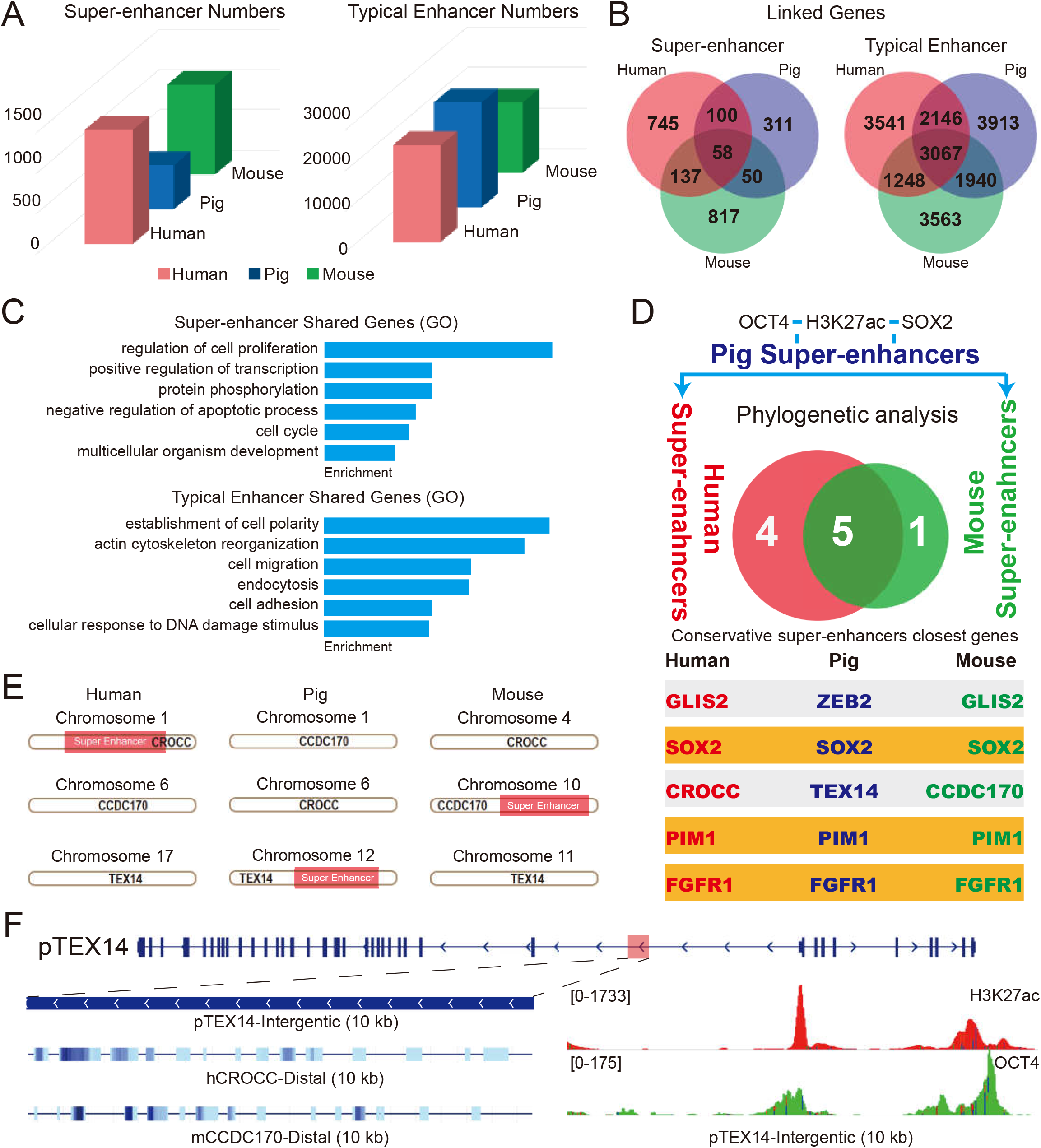
The rapid evolution of pluripotency-associated SEs in mammals. *(A)* Numbers of humans, pigs, and mice pluripotency-associated SEs and TEs. *(B)* SE- and TE-linked genes of humans (red), pigs (blue), and mice (green). *(C)* Gene Ontology (GO) analysis of the shared enhancer-linked genes in three species. *(D)* Phylogenetic analysis of the SEs sequence of humans, pigs, and mice. The H3K27ac, OCT4, and SOX2 ChIP-defined SEs in porcine PSCs were identified and subjected to conservation analysis in the genomes of humans and mice. The SE-linked genes were listed. *(E, F)* Characteristics of the SE linked to the gene TEX14(SE-TEX14) in pigs. SE-TEX14 shows different proximal genes in humans and mice, and displays high OCT4 occupation.

Next, we selected H3K27ac, OCT4, and SOX2 ChIP-defined SEs in porcine PSCs and then subjected them to conservation analysis in the genomes of humans and mice using PhastCons. There were 9 SEs with PhastCons scores larger than 0.4 between humans and pigs, and 6 between pigs and mice. Only five (ZEB2, SOX2, TEX14, PIM1, and FGFR1) conserved SEs were mapped across the three species, yet two (ZEB2 and TEX14) of them displayed discrepant proximal genes (Fig. 4D). For example, the SE linked to the gene TEX14 in pigs (SE-TEX14) is proximal to CROCC in humans and CCDC170 in mice (Fig. 4E and F). Importantly, systematic evaluation of the conservation score for SE-SOX2, SE-PIM1, and SE-FGFR1 using UCSC whole-genome alignment data (100 species) indicated that these three SEs are highly conserved in an infraclass of *Mammalia, Placentalia* (accounting for 94% of mammal species). In contrast, no or small conserved regions of the three SEs was detectable in more basal mammals (such as kangaroo and platypus), which may be due to the large phylogenetic distance or may suggest they were newly evolved in *Placentalia*. The conservation of these SEs seems to be correlated with the involvement of their target genes during early embryogenesis in humans, pigs, and mice. Altogether, our data suggest that highly conserved SEs are present in mammal genomes.

### Activation of highly conserved SEs correlates with pluripotency

We next investigated whether the activation of SE-SOX2, SE-PIM1, and SE-FGFR1 is correlated with stem cell pluripotency. Given SEs are large in size, we asked if core subregions that play a dominant role are present in the identified SEs (53). As expected, a 143 bp, 187 bp, and 124 bp subregions were identified for SE-SOX2, SE-PIM1, and SE-FGFR1, respectively, with much higher conservation scores and OCT4 occupation than the rest of the regions of each SE (Fig. 5A-C). We then performed lentivirus-based reporter assays in porcine PSCs by cloning all three subregions into a GFP reporter vector with a basal promoter which alone is not sufficient to drive the reporter expression. The expression of GFP relies on the regulatory activities of tested DNA fragments. To enhance the reporter expression, we utilized multiple tandem duplications of each core region as previously described (43). Upon the lentivirus infection, we observed robust activation of GFP for all enhancers in porcine PSCs, but not in somatic cells without pluripotency. Similarly, consistent results were also observed in mouse PSCs, suggesting the three subregions contain pluripotency-dependent enhancer activities. Because BRD4 is an essential activator for pluripotency-associated SEs, we tested if the three conserved subregions respond to BRD4 inhibition. After 48 h of 0.25 µM BRD4 inhibitor treatment, the intensity of GFP expression reduced dramatically, followed by the disappearance of GFP positive cells at 72 h post-treatment (Fig. 5D). We concluded that the three highly conserved subregions of each SE contain enhancer activities sufficient to activate pluripotency-dependent gene expression.

**Figure 5.**
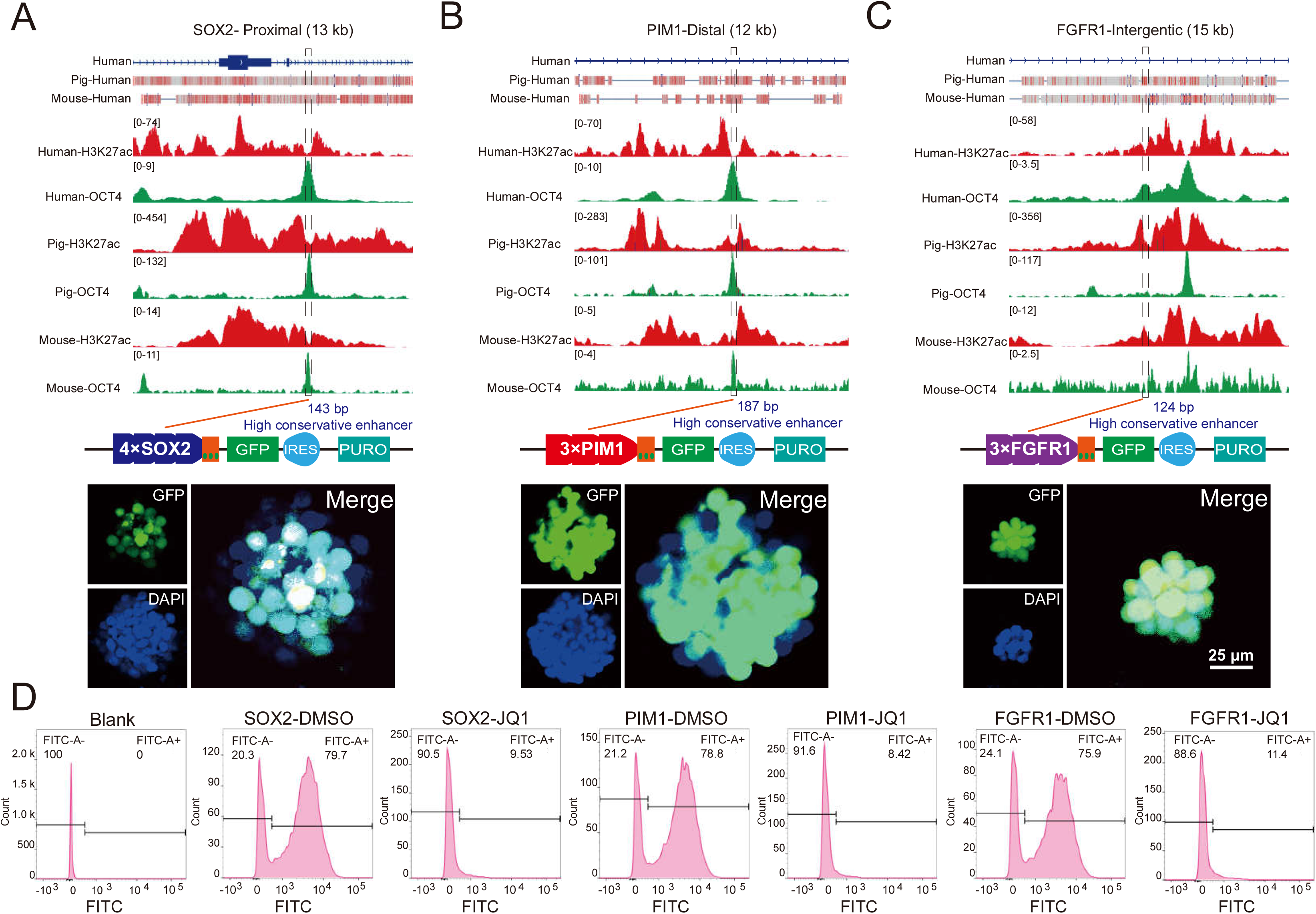
Activation of highly conserved SEs correlates with pluripotency. *(A-C)* Characteristics of the three conserved SEs. The top part is the gene models of the SEs in the human genome, and the conserved regions across species are marked by red rectangles. The middle part is the distribution of histone H3K27ac and OCT4 ChIP-seq signals in the selected SEs. The conserved subregions we identified are marked by dotted boxes. The bottom part is the results of our reporter assay. The three conserved subregions are sufficient to drive the reporter expression. The nuclei are marked by hoechst33342. Scale bar, 25 µm. *(D)* Flow cytometric analysis of GFP expression in murine PSCs. BRD4 inhibitor reduced the activity of SEs conserved subregions. The abscissa represented the GFP fluorescence intensity, and the ordinate represented the number of cells with certain GFP fluorescence intensity. The Blank group represented wild-type mouse PSCs. Cells were divided into two groups according to the GFP expression.

### Highly conserved SEs are essential for the maintenance of pluripotency

The remarkable conservation of three identified SEs in mammals indicates potential roles in controlling stem cell pluripotency (Fig. 6A). To test the function of SEs, we generated SE-disrupted mouse PSCs for SE-SOX2, SE-PIM1, and SE-FGFR1 with CRISPR/Cas9 approach by replacing each enhancer with GFP- and RFP-STOP cassettes, respectively (Fig. 6B). As expected, we observed a significant reduction in the expression of SE-linked genes in mutant clones. SE-disrupted PSCs exhibited a dramatic loss of pluripotency, spontaneous differentiation, and impaired proliferative potential, resulting in a reduced number of undifferentiated colonies compared with the control (mROSA26-locus edited) as revealed by AP staining (Fig. 6C and D). Consistent results were also observed in porcine PSCs upon the deletion of each SE using a lentivirus system expressing sgRNAs. To further verify the reduction of SE-linked gene expression in mutant clones is caused by enhancer disruption rather than a change of differentiated cell state. We decided to carry out a conditional deletion using Cre-*loxP* system. We inserted two *loxP* sites on either side of the SE-FGFR1 core subregion and knocked in a tamoxifen-inducible Cre-activation cassette at mROSA26 locus via homologous recombination. The gradual decrease of FGFR1 expression upon tamoxifen treatment at 48 and 96 hours confirmed the SE-dependent activation of target gene expression.

**Figure 6.**
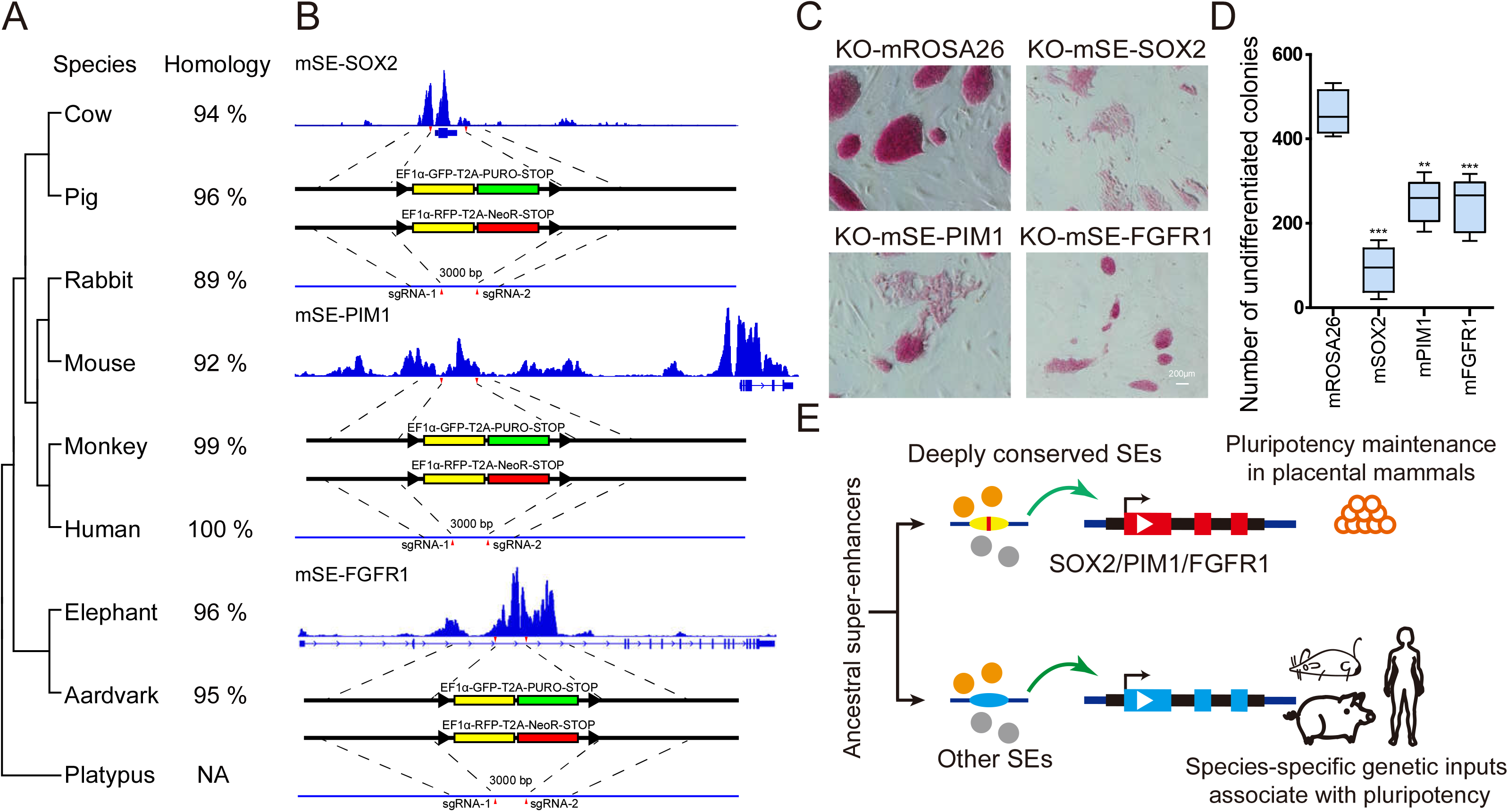
Highly conserved super-enhancers are essential for pluripotency. *(A)* The evolutionary tree of the three SEs conserved subregions among 9 representative mammals. Their sequence identities to humans were listed on the right. *(B)* The strategy to disrupt the highly conserved SEs in murine PSCs by CRISPR/Cas9. *(C)* AP staining assay of murine PSCs with SEs disruption. Murine PSCs lost their pluripotency when the three conserved SEs were destroyed. The editing of murine ROSA26 locus was used as the control. Scale bar, 200 µm. *(D)* Numbers of undifferentiated AP-positive colonies under different editing treatments. n = 5 independent experiments (\**P* < 0.05, \*\**P* <0 .01, \*\*\**P* < 0.001). *(E)* The model of the ancestral SEs evolution in mammalian pluripotency maintenance. We propose that highly conserved SEs are critical in maintaining mammalian pluripotency and other SEs may contribute to the species-specific genetic inputs associated with pluripotency.

Next, we asked whether the conserved subregions and non-conserved subregions of SEs contribute equally to the maintenance of pluripotency. We selected a non-conserved region of SE-FGFR1 and subjected it to CRISPR/Cas9-mediated deletion in mice PSCs. In contrast to conserved subregions, deletion of the non-conserved region failed to produce an obvious effect on the expression of FGFR1 and stem cell pluripotency as revealed by AP staining and cell morphology. Our results suggest that the conserved regions seem to be more important for pluripotency.

To determine whether non-conserved (species-specific) SEs also contribute to the maintenance of stem cell pluripotency, we selected a mouse-specific SE associated with the JAM2 gene as an example. We performed lentivirus-based reporter assays for SE-JAM2 and observed a similar ability to activate the reporter expression in murine PSCs compared with the three conserved SEs. Interestingly, SE-JAM2 deletion led to a significant reduction of JAM2 gene expression but has no impact on stem cell pluripotency, In addition, we observed no changes in the expression of pluripotent factors such as SOX2 upon the deletion of SE-JAM2, which further suggested that SE-JAM2 is not involved in the maintenance of pluripotency. Taken together, our data indicate that highly conserved super-enhancers are essential for maintaining stem cell pluripotency, and conserved subregions of each SE are likely to be more important than non-conserved subregions.

## Discussion

In this study, we applied comprehensive genomic and epigenomic approaches to compare pluripotency-associated SEs in pigs, mice, and humans. Although most SEs have undergone rapid changes in DNA sequence and rearrangements during mammalian evolution, we identified three highly conserved SEs linked to SOX2, PIM1, and FGFR1, respectively. The remarkable conservation of these SEs seems to be consistent with the critical roles which SE-linked genes play during development and in the maintenance of pluripotency. SOX2, PIM1, and FGFR1 are activated during early embryogenesis in humans, pigs, and mice. Further, SOX2 and FGFR1 are well-known regulators required for maintaining stem cell pluripotency (54-58). Deletion of SOX2 resulted in the trophectodermal differentiation of mouse embryonic stem cells (ESCs) (59). The expression of FGFR1 is required for the formation of prime state mouse ESCs (60) and blocking FGFR1 reduces the proliferation of human ESCs (61).

Despite PIM1 knockout mice being homozygous viable(62), inhibition of PIM1 with RNAi led to extensive differentiation of mouse ESCs after 9 days of transfection(63). In contrast, overexpression of PIM1 improved the reprogramming efficiency of mouse embryonic fibroblasts (64). It has been suggested that PIM1 plays a critical role in maintaining the telomere lengths of mouse cardiomyocytes (65). We speculated that the expression of PIM1 may facilitate the proliferative potential of stem cells by regulating the length of the telomere. Besides, PIM1 can promote cell survival rate by activating the anti-apoptotic factor BCL2 (66, 67). Stem cells with high PIM1 expression levels may overcome stage-related compatibility barriers to form interspecies chimeras effectively (33, 68). Further research is needed to elucidate the function of PIM1 in maintaining pluripotency.

The essential roles of SE-SOX2, SE-PIM1, and SE-FGFR1 have allowed us to propose that highly conserved SEs are critical to maintaining stem cell pluripotency in mammals, while the rearrangement of certain conserved SEs in different species may contribute to the variation in genetic inputs involved in regulating stem cell proliferation and differentiation. The SEs can be divided into subregions with various functions (69, 70). Several studies have reported that at least a subset of subregions is required for proper SE function, despite some subregions being redundant (17, 39, 53). Additionally, during the naive-to-primed pluripotency transition of mouse PSCs, subregions within SEs displayed pronounced differences in the dynamics of CpG methylation (71). Combined with our findings, the non-conserved domains of SEs create the possibility for evolving species-specific and lineage-specific modulation of target genes involved in pluripotency. Taken together, our study highlights the rapid evolution of SEs and certain essential pluripotency-associated SEs share a deep evolutionary origin in placental mammals. The species-specific and rearranged conserved SEs may contribute to distinct genetic inputs involved in the regulation of PSCs in different species (Fig. 6E).

## Materials and methods

### Ethics declarations

All animal experiments were approved by the Animal Care and Use Center of the Northwest A & F University and were performed in strict accordance with the Guide for the Care and Use of Laboratory Animals (Ministry of Science and Technology of the People’s Republic of China, Policy No. 2006398).

### Generation of cell lines with genomic editing

The generation of the Cre-*loxP*-mediated SE-FGFR1 conditional enhancer-deletion (3000 bp) cell line was carried out in two steps. First, the sgRNA-expressing vector targeting mROSA26 locus and the donor vector containing 1500-2500 bp homology arms and tamoxifen-induced Cre-expression cassettes (MerCreMer) were transfected into stem cells. For electric transfection, 3×10^6^ cells in 250 μL of electroporation buffer containing 15 μg plasmids were treated with 300 V for 1 ms in a 2-mm gap by BTX-ECM 830 (BTX, USA). After 12 hours, cells were washed and further incubated in the fresh medium. Further, cells were screened by 300 µg/mL G418 (Gibco, USA) for 3 days. Second, the cassette (*loxP*-enhancer-*loxP*) was introduced into the PSCs through the same strategy (transfecting one donor vector and two sgRNA-expressing vectors targeting both sides of the SE-FGFR1 core subregion). After 72 hours, limiting dilution was performed and incubation was continued for 6-8 days. Individual colonies were picked and genomic DNA was extracted for genotyping. Gene editing positive colonies were expanded for further investigation (72, 73). Tamoxifen (1 μM final concentration) was purchased from MedChem Express (Monmouth Junction, NJ, USA).

### Identification of super-enhancers

The H3K27ac-defined SEs in humans and mice PSCs were obtained from previously published research (74). The identification of H3K27ac-defined SEs in porcine PSCs was performed in four steps. First, the original SEs were identified by ROSE in porcine PSCs according to the published protocol (13, 16). The total numbers of mapped H3K27ac ChIP-seq reads per million reads were calculated and normalized by the length of each enhancer. The enhancers above the inflection point were defined as original super-enhancers. Second, we keep SEs, identified in the first step, with OCT4 and SOX2 ChIP signals. Third, SEs should be absent for H3K4me3 (marks active promoters), H3K9me3, and H3K9me2 signals in porcine PSCs. Fourth, the SE regions should be accessible with high ATAC-seq signals.

In this study, we assigned the ChIP-defined SEs to their neighboring genes, allowing a maximal distance of 100 kb between SE and the target TSS, in combined with transcriptome data, H3K4me3 (marks active promoters) ChIP-seq data, and avaiblale Hi-C data from pigs, mice, and humans (51, 52). The chromosome conformation was analyzed by HiCExplorer (75).

### Super-enhancer conservation analysis

We first identified H3K27ac, OCT4, and SOX2 ChIP-defined SEs in porcine PSCs and then subjected them to conservation analysis in the genomes of humans and mice using PhastCons (76). Eventually, we identified three SEs that are highly conserved in placental mammals. The source data used to perform the sequence alignment were downloaded from UCSC (http://genome.ucsc.edu/) (77). The evolutionary trees were measured by MEGA X (78).

### Motif enrichment analysis

Motif enrichment analysis for super-enhancers was done by Homer (version v4.10) with promoter regions as background using the vertebrate database (79). Motifs with a P-value less than 10^−12^ were considered significantly enriched (80).

## Acknowledgments

We thank Seqhealth Technology Co., LTD supplies technical support. We thank Prof. Jianyong Han (China Agricultural University) and Dr. Minglei Zhi (China Agricultural University) for their assistance in bio-technical support.

